# Aurora kinase A/AURKA interacts with the mitochondrial ATP synthase to regulate energy metabolism and cell death

**DOI:** 10.1101/2023.02.02.526754

**Authors:** Rakesh Kumar Sharma, Abderrahman Chafik, Giulia Bertolin

**Affiliations:** Univ Rennes, CNRS, IGDR (Institute of Genetics and Development of Rennes), UMR 6290, F-35000 Rennes, France

**Keywords:** metabolism, cell cycle, AURKA, mitochondrial Complex V

## Abstract

Cancer cells often hijack metabolic pathways to obtain the energy required to sustain their proliferation. Understanding the molecular mechanisms underlying cancer cell metabolism is key to fine-tune the metabolic preference of specific tumors, and potentially offer new therapeutic strategies. Here, we show that the pharmacological inhibition of mitochondrial Complex V delays the cell cycle by arresting breast cancer cell models in the G0/G1 phase. Under these conditions, the abundance of the multifunctional protein Aurora kinase A/AURKA is specifically lowered. We then demonstrate that AURKA directly interacts with the mitochondrial Complex V core subunits ATP5F1A and ATP5F1B. Altering the AURKA/ATPF1A/ATPF1B nexus is sufficient to trigger G0/G1 arrest, and this is accompanied by decreased glycolysis and mitochondrial respiration rates. Last, we discover that the roles of the AURKA/ATPF1A/ATPF1B nexus depend on the specific metabolic propensity of triple-negative breast cancer cell lines, where they correlate with cell fate. On one hand, the nexus induces G0/G1 arrest in cells relying on oxidative phosphorylation as the main source of energy. On the other hand, it allows to bypass cell cycle arrest and it triggers cell death in cells with a glycolytic metabolism. Altogether, we provide evidence that AURKA and mitochondrial Complex V subunits cooperate to maintain cell metabolism in breast cancer cells. Our work paves the way to novel anti-cancer therapies targeting the AURKA/ATPF1A/ATPF1B nexus to lower cancer cell metabolism and proliferation.

## Introduction

Breast cancer is a major contributor to all cancer cases worldwide, and a leading cause of death in women (Harbeck et al., 2019). Although the molecular origins of breast cancer are heterogeneous, this disease is accompanied by high cell proliferation and increasing metabolic and energy demands. How the cancer cell can manage its metabolic demands is a matter of intense investigation. Otto Warburg pioneered one of the first attempts to study cancer metabolism, showing that cancer cells use aerobic glycolysis over oxidative phosphorylation (Warburg, 1956). However, follow-up studies revealed that strategies to target glycolytic metabolism were only effective against cancers with this particular metabolic propensity (Pelicano et al., 2006). Over the years, a growing number of reports showed that many cancers also have high mitochondrial oxidative phosphorylation (OXPHOS) rates, indicating that they rely on mitochondrial ATP for energy supply. This is the case for epithelial cancers including breast tumors, which mainly use mitochondrial ATP to sustain their development (Whitaker-Menezes et al., 2011).

In cancer-derived cultured cells commonly used as models to explore cancer features, ATP is obtained from different metabolic pathways, including OXPHOS. This is also the case for MCF7 cells derived from breast metastatic adenocarcinoma (Guppy et al., 2002), and for AS-30D hepatoma cells (Rodríguez-Enríquez et al., 2000). Furthermore, the importance of a mitochondrial oxidative metabolism was shown to be key in B16 melanoma cells and in 4T1 breast cancer cells devoid of mitochondrial DNA (ρ0). In ρ0 cells, the re-acquisition of mitochondrial DNA enables the formation of OXPHOS complexes and restores their tumorigenesis capacity and metastatic potential like their non-ρ0 counterpart (Tan et al., 2015). Since metabolic heterogeneity appears to be contributing to drug resistance in cancer, understanding metabolic preferences and the underlying molecular mechanisms in specific cancer types is crucial for developing novel therapeutics (Pranzini et al., 2021). In this light, metabolic heterogeneity is also a feature of breast cancer cells, although the contributing factors are still not fully elucidated (Patra et al., 2021).

Since oxidative metabolism emerges as one of the critical factors in the overall metabolism of several cancer cell models, OXPHOS inhibition strategies were implemented to deepen our knowledge of the molecular mechanisms implicated. Interestingly, pharmacological strategies for OXPHOS inhibition revealed perturbations in cell cycle progression, by causing arrest at different phases of the cell cycle. Mitochondrial Complex I inhibition induced by rotenone or 1-Trichloromethyl-1,2,3,4-tetrahydro-beta-carboline (TaClo) was shown to cause G2/M arrest in a human B lymphoma cell line and in neuroblastoma cells, respectively (Armstrong et al., 2001; Sharma et al., 2017). Alternatively, Complex III inhibition by antimycin A caused S-phase arrest (Byun et al., 2008; Yh et al., 2008), while Complex V inhibition by oligomycin led to G1-phase arrest (Byun et al., 2008; Gemin et al., 2005).

As indicated above, many epithelial cancers show oxidative metabolism (Whitaker-Menezes et al., 2011), and an accompanying feature is a high expression of the cell cycle protein Aurora kinase A/AURKA (Nikonova et al., 2013). The finding that AURKA has a mitochondrial localization and that it increases mitochondrial ATP production when overexpressed provided an additional piece of evidence linking cancer biology, the cell cycle and mitochondrial metabolism (Bertolin et al., 2018). A differential level of mitochondrial AURKA was observed among various breast cancer cells used as models (Bertolin et al., 2018). However, the possibility that AURKA shapes the metabolic preference of these cells through a direct action on OXPHOS complexes has never been ruled out.

In this study, we analyzed the molecular mechanisms underlying Complex V inhibition, and their consequences on cell cycle progression and metabolic propensities in breast cancer cells. This was achieved either with a pharmacological inhibition of Complex V using oligomycin, either by downregulating core components of Complex V. While screening for cell cycle proteins with an impact on Complex V functions, we uncovered that AURKA is a physical interactor of Complex V subunits. We also showed that compromising Complex V abundance leads to profound alterations in glycolysis and oxidative phosphorylation rates of breast cancer cells. Although these cells are metabolically heterogeneous, we showed that those with high AURKA expression levels have a propensity towards an oxidative metabolism while cells with low AURKA levels have a preference for glycolysis as the main ATP source. Reducing the levels of AURKA or of its Complex V partners lowers both glycolysis and mitochondrial respiration, and it leads to G1-phase arrest and to cell death.

## Materials and Methods

### Cell culture procedures

Mycoplasma-free HEK-293 (CRL-1573) and MCF7 (HTB-22) cells were purchased from the American Type culture collection. Hs 578T and T47D were obtained as described in (Bertolin et al., 2018), and Human Primary Fibroblast were a kind gift of R. Pedeux (CLCC Eugène Marquis, Rennes, France). All cells were cultured in Dulbecco’s Modified Eagle Medium (DMEM, Thermo Fisher Scientific) supplemented with 1% penicillin-streptomycin (Thermo Fisher Scientific), and 10% FBS (Eurobio Scientific). For galactose-containing medium, DMEM glucose-free media (Thermo Fisher Scientific) was supplemented with 4.5 g/L galactose, 1% penicillin-streptomycin. Cells were cultured in 10 cm^2^ petri dishes for subcellular fractionation, and in 6-,12- or 96-well plates for all other experiments except FRET/FLIM. When cells reached 70% confluence, they were either incubated with pharmacological compounds or underwent transfection procedures. For all experiments with OXPHOS inhibitors, cells were treated with 2μM oligomycin or 0.5 μM antimycin A, and treatment durations are indicated in each corresponding Figure legend. The AURKA inhibitor MLN8237/Alisertib (Selleckchem) was used at a final concentration of 250 nM for 48 h. For control conditions, cells were stimulated with dimethyl sulfoxide (DMSO) to a final concentration of ≤0.1%. The siRNA against *AURKA* was synthesized and purchased from Eurogenetec (sequence: 5’-AUGCCCUGUCUUACUGUCA-3’), as previously described (Bertolin et al., 2016), while siRNAs against *ATP5F1A* (SI04989873), *ATP5F1B* (SI02626722), and Allstars negative control (SI03650318) were purchased from Qiagen. SiRNA transfections were done using Lipofectamine RNAiMAX reagent (Thermo Fisher Scientific) following the manufacture’s protocol. For FRET/FLIM experiments, cells were plated in four-well Nunc Lab-Tek II Chamber slides for live microscopy, and transfected with Lipofectamine 2000 according to the manufacturer’s instructions. The list of plasmids used is reported in Supplementary Table 1. Prior to imaging, cells were washed with pre-warmed, 1X Phosphate Buffered Saline (PBS) and standard growth media was replaced with phenol red-free Leibovitz’s L-15 medium (Thermo Fisher Scientific), supplemented with 20% FBS and 1% penicillin-streptomycin.

### Western blotting procedures

Total protein lysates were prepared by lysing cells in a buffer containing 50 mM Tris-HCl (pH 7.5), 150 mM NaCl, 1.5 mM MgCl2, 1% Triton X-100, and supplemented with 0.5 mM dithiothreitol (DTT), 0.2 mM Na3VO4, 4 mg/ml NaF, 5.4 mg/ml β-glycerophosphate and protease inhibitors (Complete Cocktail, Roche). Protein content was measured with the Bradford reagent (Bio-Rad), and samples were boiled in Laemmli sample buffer at 95°C for 5 min before SDS-PAGE. 20 μg of protein extracts were loaded and run on 5-20% gradient pre-cast gels (Bio-Rad) before being transferred onto a 0.45 μM nitrocellulose membrane (Amersham, Sigma-Aldrich) and blocked for one hour with 5% BSA in Tris Buffered Saline containing 0.1% Tween-20 (TBS-T). Membranes were then incubated with primary antibodies overnight at 4°C, rinsed three times with TBS-T, and incubated with species-specific secondary antibodies for one hour. They were again washed three times with TBS-T, and developed using westernBright ECL-spray (K-12049-D50, Advansta) and a CURIX 60 developer (Agfa Healthcare). Protein abundance was quantified with using Image J software (NIH).

The following primary antibodies were used: mouse anti-p53 (# 2524), rabbit anti-phospho-CDK1(Tyr15) (# 9111), rabbit anti-p27(# 3686), rabbit anti-Cyclin B1 (# 4138), rabbit anti Cyclin E1 (# 20808) and rabbit anti Cyclin C1 (# 68179), all from Cell Signaling. Similarly, rabbit anti-Cyclin D1 (Abcam, # ab134175), rabbit anti-Actin (Sigma-Aldrich, # A5060), mouse anti-AURKA (Clone 5C3, (Cremet et al., 2003)), mouse-anti ATP5FB1 (Santa Cruz, # sc-166462), mouse-anti ATP5FA1 (Invitrogen, # 459240) and mouse-anti-TOM70 (Abcam, # ab106193). The mouse anti-AURKA primary antibody was used at a 1:50 dilution, mouse anti-ATP5A and -ATP5B were used at a 1:500 dilution, and all other antibodies were used at a 1:1000 dilution. HRP-conjugated anti-mouse and anti-rabbit secondary antibodies were purchased from Jackson immunoresearch laboratories and used at a 1:10,000 dilution.

### Total ATP measurement and MTT assay

Total cellular ATP levels were measured using a bioluminescent ATP assay Kit (Promega, # G755A, G756A) as described in the manufacturer’s instructions. Luminiscence was recorded with an EnsightTM 96-well plate reader (Perkin Elmer). 3- (4,5-dimmethylthiazol-2-yl)-2,5-diphenyltetrazolium bromide (MTT) assays were used to estimate cell number following the manufacturer’s description (Tocris, # 5224), and absorbance was measured at 550 nm in an EnsightTM 96-well plate reader (Perkin Elmer).

### Colony forming assay

HEK-293 and MCF7 cells were seeded in 6-well plates at an initial seeding density of 500 cells/well. After 3-4 days, cells were treated with Complex V or Complex III inhibitors at the previously-indicated concentrations, or incubated with galactose-containing DMEM for 2-3 weeks. After each week, cells were treated with fresh media containing the inhibitors. After gently rinsing the cells with 1X PBS, cells were fixed in 4% PFA for 10 min at room temperature. Cells were then washed twice with 1X PBS and stained with 0.2% (w/v) crystal violet for 10 min. Excess of dye was removed by washing the cells with 1X PBS, before letting the wells dry at room temperature. Then, plates were scanned on Epson Perfection V700 scanner and colonies were counted using the *Colony Counter* plugin of the Image J software (NIH)

### Flow cytometry

Analysis of cell proliferation, mitochondrial membrane potential, cell cycle, cell death and LDH immunostaining were performed on a BD FACS Fortessa X20 flow cytometer. Cell proliferation was calculated using a CytoTrackTM Cell proliferation Assay Kit (Bio-Rad), and cell death was performed using Annexin V-FITC/PI apoptosis detection kit following the protocol described by the manufacturer (Bio-Rad).

Mitochondrial membrane potential and cell cycle analyses were done simultaneously by staining live cells for 30 min with Hoechst 33342 (1 μg/ml), and with TMRM (Tetramethylrhodamine methyl ester perchlorate) at a final concentration of 50 nM. After staining, a minimum of 10,000 cells were acquired for each sample. After acquisition, data were analyzed on FACS Diva software for TMRM, and cell cycle analysis was done using the Modfit LT software (Varity Software House, Tophsam). For the immunostaining of LDH, cultured cells were harvested and then fixed in 4% PFA for 10 min. Cells were then permeabilized in 0.3% Triton X-100 in 1X PBS (PBS-T) for 15 min, and then blocked in 5% BSA for 30 min. Followed by a wash in 1X PBS, cells were then incubated in a 1:100 dilution of a mouse anti-LDH primary antibody (Santa Cruz Biotechnologies, # sc133123) in PBS-T containing 1% BSA for 1 h. Cells were then washed twice and incubated with an anti-mouse secondary antibody conjugated to Alexa Fluor 488 (Thermo Fisher Scientific, # A11001) at a 1:1000 dilution in PBS-T, and supplemented with 1% BSA for 30 min. After rinsing, samples were run and results were analyzed using FACS Diva software (BD Biosciences).

### Seahorse analyses

Measurement of mitochondrial respiration was done using a Seahorse XF Cell Mito Stress Test kit (Seahorse, Agilent) as described by the manufacturer. Briefly, ~15,000 cells/well were cultured on XF24 microplates (Seahorse, Agilent). After 24 hours, cells were transfected with and incubated for further 48 hours. Then, cells were washed and incubated in XF Assay Medium supplemented with 1 mM sodium pyruvate, 2 mM glutamine and 5.5 mM D-glucose. Oxygen consumption (OCR) and extracellular acidification (ECAR) rates were measured on an XF24 extracellular flux analyzer. After plate reading, media was removed and rinsed gently with 1X PBS before cell lysis. Cells were then lysed by adding 20 μl of lysis buffer in each well for 10 min. After cell lysis, protein concentration was measured in 96 well plate format with the Bradford reagent (Bio-Rad). The OCR and ECAR values were normalized to protein content by entering the protein concentration directly in the Aglient wave 2.4 software.

### FRET/FLIM microscopy

To determine Förster’s Resonance Energy Transfer (FRET), GFP lifetime was measured as in (Demeautis et al., 2017) with an inverted SP8 Leica confocal microscope (Manheim, Germany) equipped with a Single Molecule Detection (SMD) module based on a Picoquant hardware solution (Berlin, Germany), a 470 nm pulsed laser with a 40 MHz repetition rate, a 500/50 nm band pass filter, a single-photon avalanche diode (SPAD) detector, and a 63 X oil immersion objective (N.A. =1.4). A Picoharp 300 was used for time-correlation and for image reconstruction using scanning signals to recover Fluorescence Lifetime Imaging Microscopy (FLIM) images (time-tagged time resolved (TTTR) method). Lifetime values in each individual cell were determined by fitting the fluorescence decay extracted. This was performed by integrating the signal from the pixels from a Region Of Interest (ROI) with a single exponential model. Fitting was performed with the Symphotime software.

### Statistical analyses

Two-way ANOVA tests were performed to compare two variables among multiple conditions, one-way ANOVA tests were used when one variable was tested among multiple conditions, and Student’s t-tests were used to compare two conditions. Data were checked for normality before performing any statistical test.

Two-way ANOVA and the Holm-Sidak method were used to compare the effect of each OXPHOS inhibitor and of the cell cycle phase on the percentage of cells in each cell cycle phase (Fig. 1B and Supplementary Figs. 1D, 2B), and the effect of oligomycin and of time on Cytotrack fluorescence intensity (Fig. 1F and Supplementary Fig. 1G). One-way ANOVA and Dunnett’s multiple comparison test were employed to compare the effect of oligomycin and antimycin A on the relative abundance of cell cycle-related proteins (Fig. 2A and Supplementary Fig. 3), the effect of the cell line model on: OCR/ECAR ratios (Supplementary Fig. 5B), and on the percentage of cells with high LDH (Supplementary Fig. 5D), and the effect of siRNAs on: the percentage of cells with low TMRM fluorescence (Fig. 3A), on the percentage of cells in G0/G1 (Fig. 3B, F, G), on total ATP levels (Fig. 3C, F), on OCR (Fig. 4A, D, G) and on ECAR (Fig. 4B, E, H) rates and on PI/annexin staining (Fig. 5A, B). One-way ANOVA on ranks and Dunn’s method were used to compare ΔLifetime values (Fig. 2B, C), the effect of siRNAs on total ATP levels (Fig. 3G), and the effect of oligomycin and antimycin A on the relative abundance of cell cycle related proteins (Supplementary Fig. 3).

**Figure 1.**
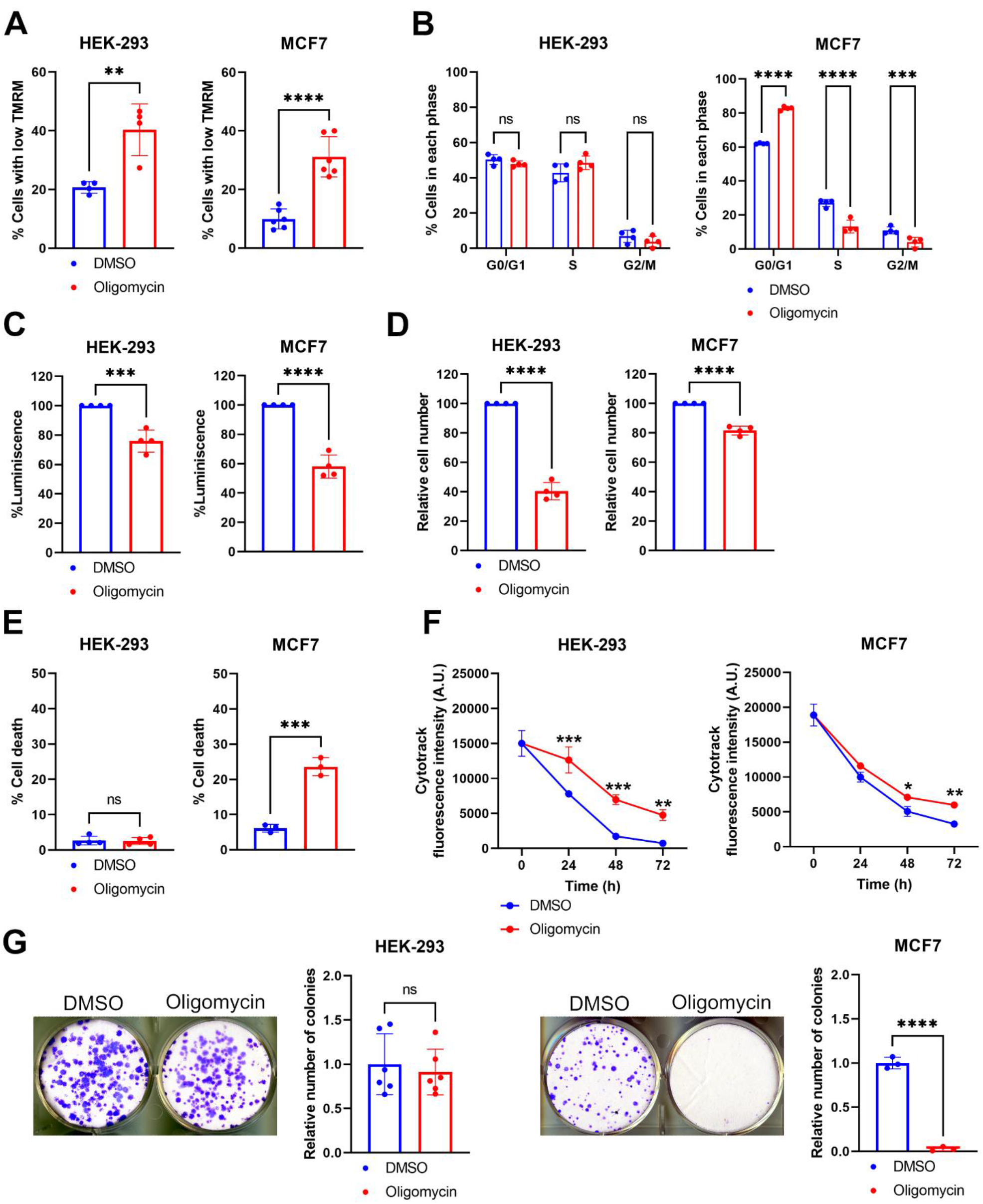
Mitochondrial Complex V inhibition with oligomycin alters cell cycle progression in MCF7 cells, but not in HEK-293 cells. (**A**) Percentage of cells with low TMRM fluorescence intensity in HEK-293 (left) and MCF7 (right) cells treated with DMSO or oligomycin for 48 h. (**B**) Percentage of HEK-293 (right) or MCF7 cells stained with the nuclear dye Hoechst 33324 to identify the phases of cell cycle, and following an incubation with DMSO or oligomycin for 48 h. (**C**) Percentage of total ATP levels in HEK-293 (left) or MCF7 (right) cells treated with DMSO or oligomycin for 48 h. ATP levels were measured using a luminescence-based assay, and were relative to the DMSO condition. (**D**) Relative HEK-293 (left) or MCF7 (right) cell number upon treatment with DMSO or oligomycin for 72 h, measured with an MTT assay. (**E**) Proportion of HEK-293 (left) or MCF7 (right) cells showing cell death features identified with PI/annexin stainings after treatment with DMSO or oligomycin for 72 h. (**F**) Cytotrack mean fluorescence intensity in HEK-293 (left) or MCF7 (right) cells treated with DMSO or oligomycin for the indicated timepoints. (**G**) Representative images and corresponding quantifications of colony-forming assays in HEK-293 (left) or MCF7 (right) cells, treated with DMSO or oligomycin for at least 2 weeks. The number of colonies in the oligomycin-treated condition is relative to that of each corresponding control. Data are means ± SD. \**P*<0.05,\*\**P*<0.01, \*\*\**P*<0.001, and \*\*\*\**P*<0.0001 compared to each corresponding DMSO condition (A-E, G), or to each corresponding time point in the DMSO condition (F). ns: not significant. *n* ≥ 3 independent experiments with at least 10,000 cells per condition quantified in A, B, E, F. *n* ≥ 3 independent experiments for all other panels. A. U.: arbitrary units.

Student’s t-test was used to compare the percentages of cells with low TMRM intensities (Fig. 1A and Supplementary Fig. 1C), total ATP levels (Figs. 1C, 3D and Supplementary Figs. 1E, 2A, 5E), cell number (Fig. 1D), cell death features (Fig.1E and Supplementary Figs. 1F, 2C, 6), the relative number of colonies (Fig. 1G and Supplementary Fig. 2D), the percentage of cells in G0/G1 (Fig. 3D, E). The Kolgorov-Smirnoff (Fig. 3E) or the Mann-Whitney tests (Supplementary Fig. 2A) were also used to compare total ATP levels. Alpha for all statistical tests was equal to 0.05.

## Results

### Mitochondrial dysfunction by Complex V inhibition reveals a cell cycle delay specific to breast cancer cells

It has previously been shown that treatment with mitochondrial respiratory complex inhibitors causes cell cycle arrest in specific cell models. For instance, incubating HeLa-derived Chang cells or IL-60 cells with the mitochondrial Complex V inhibitor oligomycin induces G1 arrest (Gemin et al., 2005; Byun et al., 2008). After oligomycin treatment, Th17 immune cells also show significant cell proliferation defects, which may support a cell cycle delay or arrest in non-cancerous cell lines as well (Shin et al., 2020). However, it is unclear whether these effect are limited to specific cancer cell lines, or whether they are shared by non-cancerous cells as well. To answer this question, we screened the effect of oligomycin in three independent cell models: MCF7 human breast cancer cells, non-tumorigenic HEK-293 Human Embryonic Kidney cells, and Human Primary Fibroblasts (HPFB). All cell models were treated with oligomycin for 48 h, and we explored the consequences of such treatment on mitochondria- and cell cycle-related parameters such as mitochondrial membrane potential, cell cycle progression, ATP production, cell proliferation and cell death.

Mitochondrial membrane potential and cell cycle analyses were done simultaneously in live cells co-stained with the mitochondrial potentiometric dye tetramethylrhodamine, methyl ester (TMRM) and with the nuclear dye Hoechst 33342, followed by flow cytometry analyses. Oligomycin caused a significant increase in the number of cells with low TMRM intensity in HEK-293 and in MCF7 cells compared to their DMSO-treated counterparts (Fig. 1A, Supplementary Fig. 1A). This indicates that long-term treatment with oligomycin induces mitochondrial depolarization. While short-term treatments with this compound are known to induce an increase of the mitochondrial membrane potential (ΔΨ) (Kalbáčová et al., 2003; Macouillard-Poulletier de Gannes et al., 1998; Yang et al., 2021), our data are in line with previous reports showing that long incubations with oligomycin ultimately lead to ΔΨ loss (Kalbáčová et al., 2003; Yang et al., 2021). Under the same treatment conditions, we also monitored the incorporation of Hoechst 33342 and we observed an alteration in cell cycle progression specific to MCF7 cells only. Compared to controls, the amount of cells in G0/G1 increased by 20% upon oligomycin treatment, whereas cells in S and G2/M phases decreased by 13% and 6%, respectively (Fig. 1B, Supplementary Fig. 1B). Conversely, we didn’t see any comparable difference in the cell cycle profiles of HEK-293 treated with DMSO or with oligomycin (Fig. 1B, Supplementary Fig. 1B).

Next, we evaluated the impact of oligomycin on total ATP production using a luciferase-based ATP estimation assay. Oligomycin treatment in non-tumorigenic HEK-293 cells induced a 24% reduction in total ATP compared to cells treated with DMSO, and a reduction of ~ 42% in MCF7 cancer cells (Fig. 1C). We also tested the impact of oligomycin on non-transformed HPFB. While we observed a significant difference on the percentage of cells with low TMRM after oligomycin treatment, no changes in cell cycle progression and in total ATP levels were observed compared to control cells (Supplementary Fig. 1C-E). This indicates that in this cell model, lowering mitochondrial membrane potential is not sufficient to induce an arrest in the G0/G1 phase, nor it triggers an overall loss of total ATP levels.

Since we were intrigued by the G0/G1 cell cycle arrest and the dramatic ATP decrease induced by oligomycin in MCF7 cells, we asked whether these effects were related to increased cell death events or lower proliferation rates. To this end, we first compared the consequences of oligomycin on 3-(4,5-dimmethylthiazol-2-yl)-2,5-diphenyltetrazolium bromide (MTT), which is considered as a readout of the overall cell number (Mosmann, 1983; Stockert et al., 2018), in MCF7 and in HEK-293 cells. HEK-293 cells showed a drastic reduction in cell number, with only 40% of the cells effectively reducing MTT after a 72 h-long oligomycin treatment (Fig. 1D). To verify whether this reduction in cell number was due to an increase in cell death events or to reduced cell proliferation, we stained the cells with Propidium Iodide (PI) and annexin V and we analyzed them by flow cytometry (Vermes et al., 1995). HEK-293 showed no signs of ongoing cell death events – presented as the sum of early apoptotic, late apoptotic, and necrotic stages – (Fig. 1E). Similarly, oligomycin did not induce any increase in cell death events in HPFB (Supplementary Fig. 1F). This supports the possibility that the loss of ATP and MTT activity observed in HEK-293 cells is due to reduced cell proliferation rates. On the contrary, the great majority (~80%) of MCF7 cells were able to reduce MTT upon incubation with oligomycin (Fig. 1D). However, oligomycin significantly increased the amount of cell death events in MCF7 cells, again pointing at cell-specific differences when cells are treated with this compound (Fig. 1E).

To obtain an insight on cell proliferation rates, we measured the effect of oligomycin on the readout of the Cytotrack Green cell proliferation assay. Cytotrack is a cell permeable dye containing a fluorophore and a fluorescence blocker. Upon incubation in live cells, the fluorescence blocker is cleaved by the intracellular esterases and the dye is then capable of emitting fluorescence. Such fluorescence is halved after each successive cell division, and Cytotrack fluorescence intensity loss over time can then be used as a proxy to monitor cell proliferation. To explore the effect of Complex V inhibition on cell proliferation, cells were incubated with oligomycin and harvested at three time points after treatment – 24 h, 48 h and 72 h –. Fluorescence intensity for each time point was then measured by flow cytometry. We found a significant difference in the mean fluorescence intensity of Cytotrack between DMSO- and oligomycin-treated cells in HEK-293 at all time points (Fig. 1F). Differences in Cytotrack intensity over time were less prominent for MCF7 cells (Fig. 1F) and were not significant for HPFB (Supplementary Fig. 1G), further indicating that oligomycin affects cell proliferation predominantly in HEK-293 cells and that the loss in the capacity of cells to reduce MTT could be related to a loss of cell proliferation rates. These findings were complemented by colony forming assays to determine long-term proliferation and survival. In these experiments, 500 cells were seeded and the number of colonies were counted after three weeks of treatment. We observed that oligomycin completely inhibited colony formation in MCF7 cells, but not in HEK-293 (Fig. 1G). Again, these data indicate that oligomycin treatment mainly induces cell death in MCF7 cells, while it decreases proliferation in HEK-293 cells.

Overall, our results show that impairing mitochondrial activity with oligomycin has different effects according to the cell type. In non-proliferative cells as HPFB, it increases the number of cells showing mitochondrial depolarization, but without any further effect on ATP production, cell cycle progression, overall cell number or proliferation rates. In highly-proliferative, non-tumorigenic cells as HEK-293, mitochondrial depolarization is accompanied with a mild lowering of ATP levels and a delay in proliferation rates. In highly proliferative and tumorigenic cells as MCF7 instead, oligomycin-induced mitochondrial depolarization correlates with a dramatic ATP loss, a cell cycle arrest in G0/G1 and significant cell death rates.

### Cell cycle arrest in G0/G1 is specific to Complex V inhibition in MCF7 cells

Since Complex V inhibition showed a G0/G1 cell cycle arrest along with ATP loss and increased cell death specifically in MCF7 cells, we asked whether similar effects could be achieved using alternative OXPHOS inhibitors. To this end, we inhibited Complex III for 48 h with antimycin A and we monitored its effect on the total ATP production, cell cycle profiles and cell death rates. Of note, antimycin A was previously reported to induce an early S-phase arrest in Chang cells (Byun et al., 2008). Therefore, we compared all thee cell models to verify whether the effect of antimycin A was specific to MCF7, or whether it was shared across the three cell lines. HEK-293 cells treated with antimycin A showed a loss in more than 50% of total ATP levels, and similar results were obtained in MCF7 cells (Supplementary Fig. 2A). On the contrary, no significant loss in ATP was observed in HPFB. When we looked at cell cycle profiles, a cell cycle delay with an increased number of cells in S phase was detected in the three cell types (Supplementary Fig. 2B). All three cell lines also showed an increase in the number of cell death events (Supplementary Fig. 2C). Last, colony forming assays revealed that treating cells with antimycin A induced a dramatic decrease in long-term proliferation rates of MCF7 and HEK-293 cells (Supplementary Fig. 2D).

Taken together, we confirm and complement previous data reporting that antimycin A induces an S-phase arrest in three independent cell types (Gemin et al., 2005; Byun et al., 2008), which is accompanied with cell proliferation defects and cell death events. Although blocking OXPHOS complexes globally induces a cell cycle arrest, our data substantiate previous findings indicating that the specific phase of arrest depends on which complex is inhibited (Byun et al., 2008).

### Complex V inhibition acts on the cell cycle by reducing the abundance of Aurora kinase A/AURKA

Given that the effect of mitochondrial Complex V inhibition with oligomycin specifically induces G0/G1 arrest in breast cancer cells, we sought to investigate the molecular mechanisms involved. To this end, we screened the abundance of various cell cycle-related proteins in MCF7 cells using western blotting approaches. Cells were treated for 48 h either with DMSO, with oligomycin to induce Complex V inhibition, or with antimycin A to trigger Complex III inhibition. Given that antimycin A induces S-phase arrest irrespectively of the cell line analyzed, we used this compound as a control for the specificity of changes in protein abundance observed with oligomycin.

We did not detect any significant difference in the abundance of cell cycle-related proteins known to play roles in the early phases of the cell cycle, such as CCNC (Cyclin C), CCNE1 (Cyclin E1), the G1 marker p27 or in the tumor suppressor protein p53 (Supplementary Fig. 3). Conversely, we found that the abundance of Aurora kinase A/AURKA and that of CCND1 (Cyclin D1) was significantly lowered in cells treated with oligomycin, although we found a significant decrease of CCND1 levels also in antimycin A-treated cells (Fig. 2A). Since CCND1 abundance was lowered regardless of the treatment performed, we focused on AURKA as it showed a decreased abundance specifically in oligomycin-treated conditions. In addition to the kinase, we also explored the abundance of two key cell cycle-related molecular partners of AURKA, such as CDK1 and CCNB1 (Cyclin B1). CDK1 and CCNB1 are known to associate in a complex important for mitotic initiation (Jackman et al., 1995), and CDK1 undergoes phosphorylation on Tyr15 at mitotic entry (De Souza et al., 2000). In addition, AURKA is involved in the mitotic activation of CCNB1 at the mRNA and at the protein levels (reviewed in (Nikonova et al., 2013), while CDK1 was shown to directly regulate AURKA activation and activity during meiosis in *Xenopus* oocytes (Maton et al., 2003). Last, AURKA, CDK1 and CCNB1 cooperate in the regulation of mitochondrial segregation at mitosis (Kashatus et al., 2011). Although CDK1 and CCNB1 showed a tendency towards a decrease in oligomycin-treated cells, we did not find any significant difference in the abundance of neither CCNB1, nor of CDK1 pTyr15 (Supplementary Fig. 3). This suggests that the decrease of AURKA levels upon Complex V inhibition is partially decorrelated from that of at least two of its mitotic partners.

**Figure 2.**
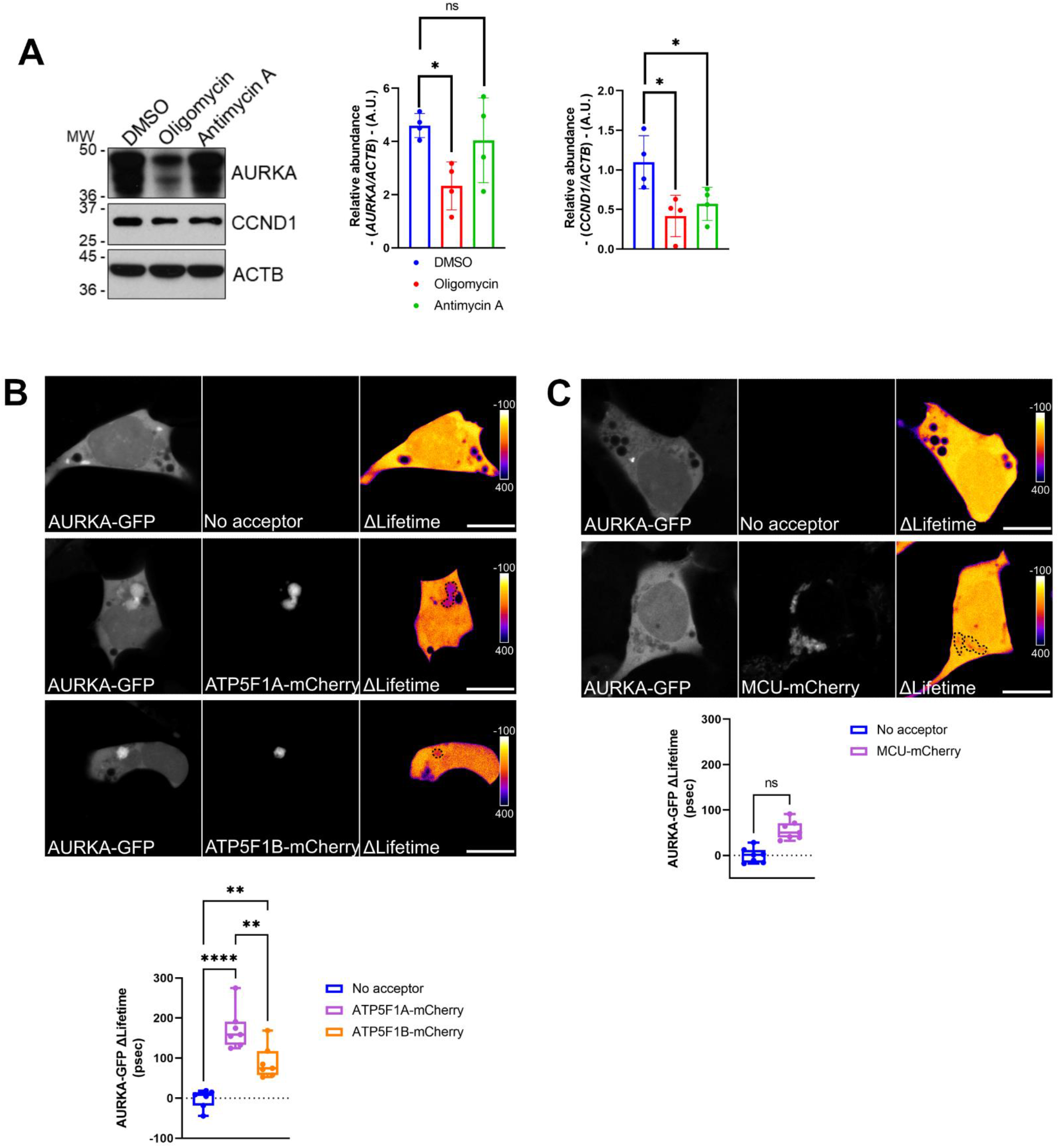
The cell cycle-related protein AURKA interacts with core Complex V subunits, and its abundance is lowered after Complex V inhibition. (**A**) Representative western blot and corresponding quantification of the abundance of the cell cycle-related proteins AURKA and CCND1 in total lysates of MCF7 cells treated with oligomycin or Antimycin A for 48 h. Loading control: ACTB. Data are from *n* = 4 independent experiments, and presented as means ± SD. A. U.: arbitrary units. (**B**, **C**) FRET/FLIM analyses on MCF7 cells expressing (**B**) AURKA-GFP alone or together with ATP5F1A-mCherry or ATP5F1B-mCherry, or (**C**) AURKA-GFP alone or together with MCU-mCherry. Scale bar: 10 μm. Pseudocolor scale: pixel-by-pixel Δlifetime. Graphs: ΔLifetime quantifications in the dotted areas. Data extend from minimum to maximum. *n* = 7 cells issued from one representative experiment (of three). \**P*≤0.05,\*\**P*≤0.01, and **** *P*≤0.0001 compared to each corresponding control condition. ns: not significant.

Given the effect of Complex V inhibition on the abundance of AURKA, we asked whether Complex V subunits could be direct interactors of the kinase. This possibility was already supported by previous mass spectrometry data (Bertolin et al., 2018). Therefore, we performed Förster’s Resonance Energy Transfer (FRET)/Fluorescence Lifetime Imaging Microscopy (FLIM) analyses to visualize the physical proximity between AURKA and two core Complex V subunits, ATP5F1A and ATP5F1B. These proteins were previously identified among the main interactors of the kinase at mitochondria (Bertolin et al., 2018). FRET/FLIM is a useful technique to detect protein-protein proximities occurring at a distance of less than 10 nm, which is compatible with molecular interactions. At the visual level, FRET can be represented with a ΔLifetime-based pixel by pixel cartography (Bertolin et al., 2021). A positive ΔLifetime was detected in subcellular areas where AURKA-GFP and ATP5F1A-mCherry or ATP5F1B-mCherry colocalized (Fig. 2C), indicating protein-protein proximities between AURKA and the Complex V subunits tested. We also noticed a profound aggregation of mitochondria in the areas where AURKA, ATP5F1A or ATP5F1B are capable of performing FRET (Fig. 2B). This was previously observed in conditions where AURKA is capable of triggering mitochondrial turnover by mitophagy, and where the organelles spared from degradation have the tendency to aggregate (Bertolin et al., 2021). To further exclude that this proximities were due to the tight environment of the Inner Mitochondrial Membrane (IMM), we calculated ΔLifetime variations between AURKA-GFP and the IMM protein Mitochondrial Calcium Uniporter (MCU), again fused to the mCherry acceptor. While mitochondrial aggregation was still observable, no significant FRET was detected between AURKA and MCU (Fig. 2C). Therefore, this corroborates the specificity of the protein proximities between AURKA and ATP5F1A and ATP5F1B.

Therefore, our data show that oligomycin treatment specifically lowers the abundance of AURKA. This might be a direct effect of Complex V inhibition, as AURKA is capable of showing protein-protein proximities compatible with molecular interactions with Complex V core subunits.

### Downregulation of AURKA, ATP5F1A or ATP5F1B phenocopies Complex V inhibition by triggering cell cycle arrest in G0/G1 and ATP loss

After observing that AURKA is at a FRET-compatible distance with ATP5F1A and ATP5F1B, we asked whether these proteins could be a nexus regulating cell cycle progression and ATP levels in breast cancer cells. To this end, we downregulated each of the three proteins using an siRNA-mediated strategy. After 48 h of depletion, MCF7 cells were co-stained with TMRM and Hoechst 33342, and analyzed by flow cytometry. Similarly to what observed in oligomycin-treated cells (Fig. 1), *AURKA* downregulation (Supplementary Fig. 4A) increased the amount of cells showing a lower mitochondrial membrane potential (Fig. 3A). Conversely, no difference was observed in cells depleted for *ATP5F1A* or *ATP5F1B* when compared to control cells (Fig. 3A, Supplementary Fig. 4A). Despite this results, we observed that *AURKA, ATP5F1A* or *ATP5F1B* downregulation was capable of increasing the number of cells found in the G0/G1 phase (Fig. 3A, B). Likewise, the downregulation of any of the three proteins led to a substantial decrease in ATP levels (Fig. 3C).

**Figure 3.**
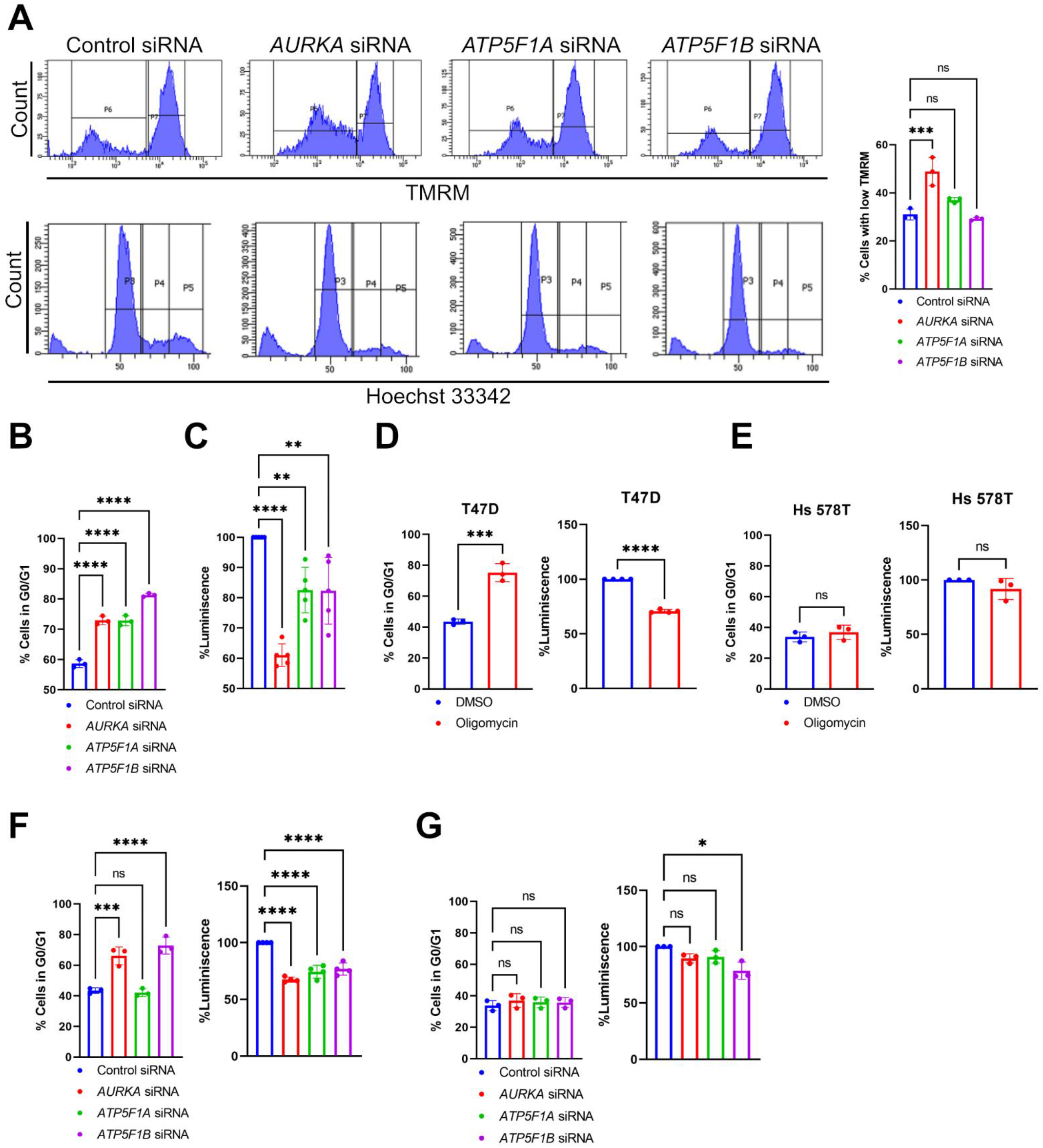
Downregulation of *AURKA, ATP5F1A* or *ATP5F1B* induces a cell cycle arrest in G1 and lowers ATP production. (**A**) Representative FACS plots illustrating TMRM fluorescence intensity (upper panels) and Hoechst 33324 fluorescence intensity (lower panels) to identify the stages of cell cycle in MCF7 cells transfected as indicated. Graph: quantification of the percentage of cells with low TMRM fluorescence intensity. (**B**) Quantification of the percentage of MCF7 cells in the G0/G1 phase of the cell cycle quantified after Hoechst 33324 incorporation, and transfected as indicated. (**C**) Percentage of total ATP levels in MCF7 cells transfected with a control siRNA, or with siRNAs targeting *AURKA, ATP5F1A* or *ATP5F1B.* ATP levels were measured with a luminescence-based assay, and were relative to the control siRNA condition. (**D-E**) Quantification of the percentage of T47D (**D**) or of Hs 578T (**E**) cells in the G0/G1 phase of the cell cycle after Hoechst 33324 incorporation (left panels), and percentage of total ATP levels (right panels) in cells treated with DMSO or oligomycin for 48 h. (**F-G**) Quantification of the percentage of T47D (**F**) or of Hs 578T (**G**) cells in the G0/G1 phase of the cell cycle after Hoechst 33324 incorporation (left panels), and percentage of total ATP levels (right panels) in cells transfected as indicated. Data are means ± SD. \**P*<0.05,\*\**P*<0.01, \*\*\**P*<0.001, and \*\*\*\**P*<0.0001 compared to each corresponding control. ns: not significant. *n* ≥ 3 independent experiments with at least 10,000 cells per condition quantified.

Thus, the silencing of *AURKA, ATP5F1A* or *ATP5F1B* recapitulates the cell cycle delay and total ATP loss previously observed in cells treated with oligomycin. Furthermore, it supports the notion that AURKA cooperates with core Complex V subunits to maintain cell cycle progression and the overall cellular energetic capacity.

To establish a putative link between GO0/G1 arrest, loss in ATP levels and the abundance of AURKA, we asked whether cell cycle progression and ATP levels are enhanced in breast cancer cells with high levels of AURKA, and whether they are reduced in breast cancer cells with low levels of the kinase. To this end, we used T47D and Hs 578T triple-negative breast cancer cells. These models were previously shown to have a high and a low abundance of AURKA, respectively, both at the total and at the mitochondrial level (Bertolin et al., 2018). In these cells, we first explored whether oligomycin treatment induced a cell cycle arrest in G0/G1, and a loss of ATP levels. While a pharmacological inhibition of Complex V could recapitulate both events in T47D cells expressing high levels of AURKA (Fig. 3D), no cell cycle arrest and no differences in ATP levels could be retrieved in Hs 578T cells (Fig. 3E). To verify whether the molecular mechanisms regulating cell cycle delay and lowered ATP levels in T47D are the same as in MCF7 cells, we downregulated *AURKA, ATP5F1A* or *ATP5F1B* (Supplementary Fig. 4B). In this model, a higher proportion of cells in G0/G1 was observed when *AURKA* or *ATP5F1B* were depleted (Fig. 3F). Although *ATP5F1A* downregulation did not result in cell cycle delay, the depletion of all three proteins induced a loss in total ATP (Fig. 3F). Conversely, *AURKA, ATP5F1A* or *ATP5F1B* silencing did not induce a G0/G1 arrest in Hs 578T cells, and only minor differences in ATP levels were detected in this model when ATP5F1B was depleted (Fig. 3G, Supplementary Fig. 4C).

Altogether, these results indicate that AURKA, ATP5F1A and ATP5F1B cooperate in regulating cell cycle progression and in maintaining total ATP levels in breast cancer cells with high AURKA expression.

### Depletion of AURKA, ATP5F1A or ATP5F1B impairs the respiratory and glycolytic metabolism of breast cancer cells

We showed here above that *AURKA, ATP5F1A* or *ATP5F1B* silencing recapitulates the main features of Complex V inhibition, including lowered ATP rates. Therefore, we asked whether this altered energetic capacity was due to an impairment of glycolysis or of the mitochondrial respiratory chain efficiency. Taking advantage of a Seahorse XF analyzer, we explored Oxygen Consumption Rates (OCR) and Extracellular Acidification Rates (ECAR) of MCF7, T47D and Hs 578T cells depleted or not of these proteins. These are a readout of the efficiency of OXPHOS and of glycolysis, respectively.

First, we evaluated the basal metabolism of the three cell types. We noticed that MCF7 (Fig. 4A, C) and T47D cells (Fig. 4D, F) transfected with a control siRNA were showing higher OCR rates than Hs 578T cells (Fig. 4G, I), while ECAR rates were slightly higher in Hs 578T cells than in the other cell types (Fig. 4B-C, E-F, H-I). We then sought to ascertain that Hs 578T cells relied more on glycolysis than MCF7 or T47D cells. When comparing the metabolic profiles of all cell types, Hs 578T cells had higher ECAR rates than MCF7 or T47D cells (Supplementary Fig. 5A, B), a higher proportion of Hs 578T cells expressed the glycolytic marker Lactate Dehydrogenase (LDH) (Supplementary Fig. 5C, D) and showed defects in total ATP rates and long-term cell proliferation in culture media containing galactose as a carbon source (Supplementary Fig. 5E, F). Once validated the basal metabolism of these cell models, we evaluated OCR and ECAR rates upon the downregulation of *AURKA, ATP5F1A* or of *ATP5F1B.* Under these conditions, all cell lines showed a dramatic decrease of both ATP sources (Fig. 4A-I). Given that ECAR analyses were performed in the presence of known OCR stressors, these results also show that altering OCR impacts ECAR as a consequence.

**Figure 4.**
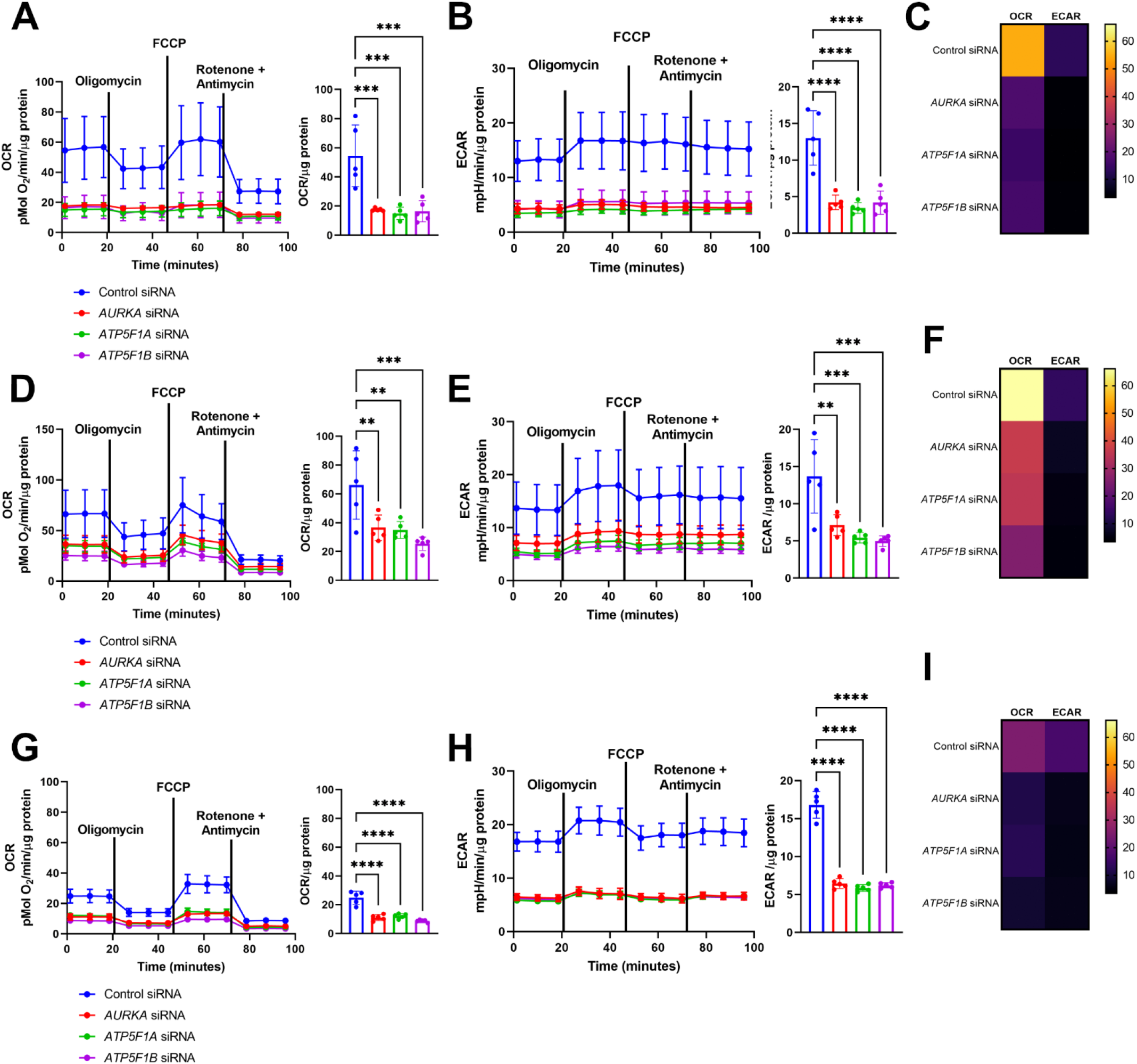
Downregulation of *AURKA, ATP5F1A* or *ATP5F1B* lowers cell metabolism by reducing oxygen consumption (OCR) and extracellular acidification (ECAR) rates. (**A**) Time-resolved kinetic graph (left) and maximal OCR capacity (right) of MCF7 cells transfected with a control siRNA, or siRNAs targeting *AURKA, ATP5F1A* or *ATP5F1B* and measured using the Seahorse technology. 2.5 μM oligomycin was used to determine ATP-linked respiration, 1μM FCCP to determine maximal respiration, and the 0.5 μM rotenone/antimycin A mix was used to determine mitochondrial spare capacity. (**B**) Time-resolved kinetic graph (left) and maximal ECAR capacity (right) of MCF7 cells transfected and treated as indicated, and measured using the Seahorse technology. (**C**) Energy heat map of maximal OCR and ECAR rates in MCF7 cells transfected as indicated. (**D, G**) Time-resolved kinetic graph (left) and maximal OCR capacity (right) of T47D (**D**) or Hs 578T (**G**) cells transfected, treated and measured as in (**A**). (**E, H**) Time-resolved kinetic graph (left) and maximal ECAR capacity (right) of T47D (**E**) or Hs 578T (**H**) cells transfected, treated and measured as in (**B**). (**F, I**) Energy heat maps of maximal OCR and ECAR rates in T47D (**F**), or Hs 578T (**I**) cells transfected as indicated. Pseudocolor scale: high-to-low OCR and ECAR maximal values. Data are means ± SD. \**P*<0.05,\*\**P*<0.01, \*\*\**P*<0.001, and \*\*\*\**P*<0.0001 compared to each corresponding control. *n* ≥ 3 independent experiments.

Altogether, these data further support the direct role of AURKA, ATP5F1A and ATP5F1B within a potential metabolic nexus in triple-negative breast cancer cells. Impairing this nexus alters both glycolytic and oxidative metabolism rates, and this regardless of the specific metabolic propensities of the cells analyzed.

### The AURKA/ATP5F1A/ATP5F1B nexus differentially regulates cell cycle arrest or cell death in cells with opposite metabolic propensities

Our Seahorse analyses illustrated the metabolic consequences of impairing Complex V functionality in cells heavily relying on mitochondria, such as T47D and MCF7 cells. Ultimately, these alterations are concomitant with a cell cycle arrest in G0/G1 (Fig. 3B, D, F). Despite observing similar OCR and ECAR variations in cells with a glycolytic propensity such as Hs 578T cells, we were surprised to see that impairing Complex V functionality in this model had no consequences on cell cycle progression (Fig. 3E, G). Therefore, we asked whether this could be explained by a differential capacity of Hs 578T cells to overcome a cell cycle arrest, and to undergo cell death events upon Complex V impairment at higher rates than cells with an oxidative metabolism. To verify whether glycolytic Hs 578T cells had a capacity to undergo cell death different from the one of oxidative T47D cells, we compared their propidium iodide (PI)/annexin profiles upon *AURKA, ATP5F1A* or of *ATP5F1B* silencing. *AURKA* and *ATP5F1B* downregulation significantly increased cell death rates in Hs 578T cells, while no effect was seen when *ATP5F1A* was silenced (Fig. 5A). A similar effect was achieved when AURKA was inhibited with the ATP analog MLN8237, a compound preventing AURKA activation and activity towards its substrates (Damodaran et al., 2017) (Supplementary Fig. 6). Conversely, no significant cell death was observed in T47D cells in any condition tested (Fig. 5B).

**Figure 5.**
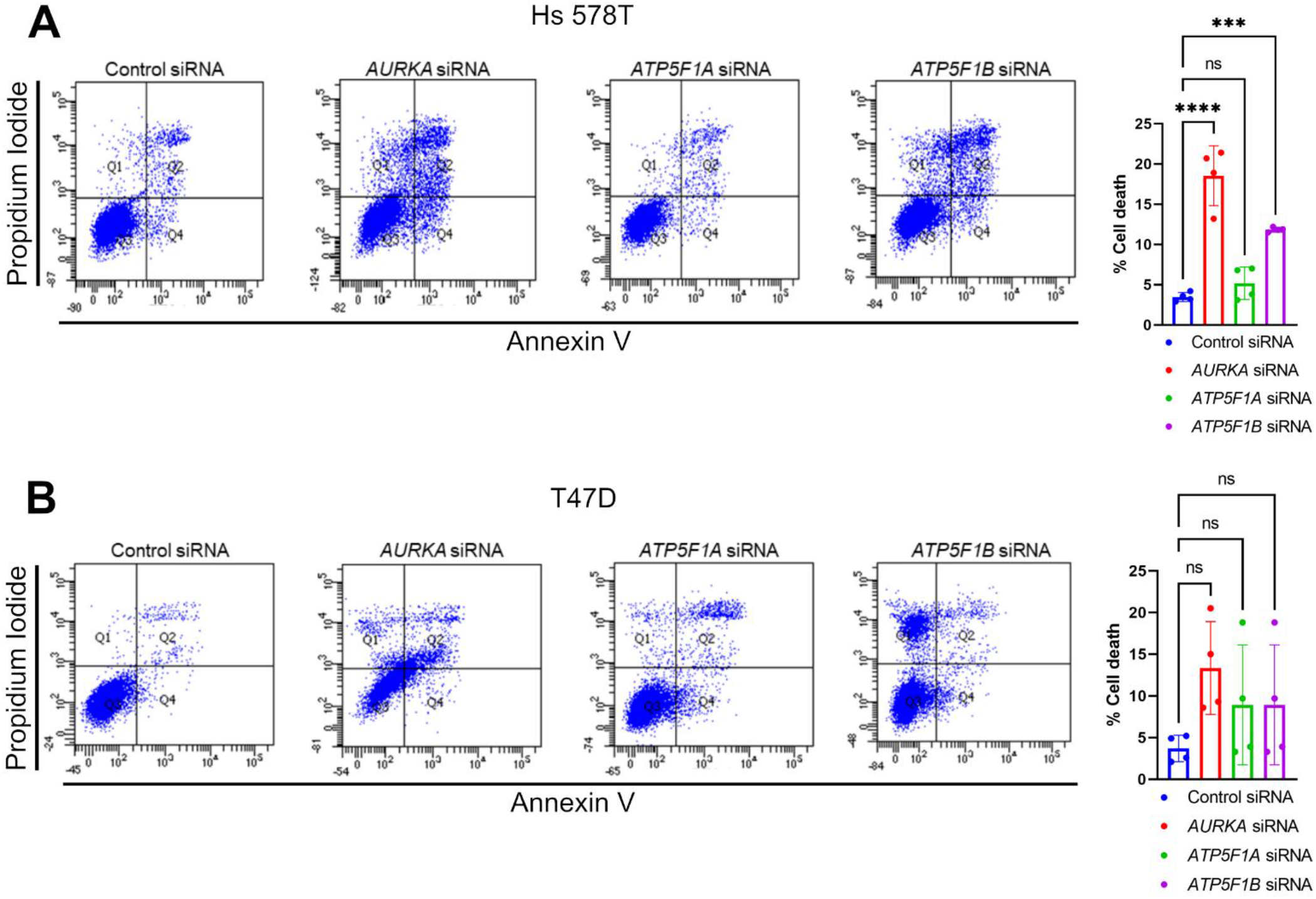
Downregulation of *AURKA, ATP5F1A* or *ATP5F1B* triggers cell death events in Hs 578T cells. (**A-B**) Representative FACS plots and corresponding quantification (Q2 section) of the percentage of Hs 578T (**A**) or T47D (**B**) cells showing PI/annexin stainings upon transfection with a control siRNA, or siRNAs targeting *AURKA, ATP5F1A* or *ATP5F1B.* Data are means ± SD. \*\*\**P*<0.001 and \*\*\*\**P*<0.0001 compared to each corresponding control siRNA condition. ns: not significant. *n* ≥ 3 independent experiments with at least 10,000 cells per condition quantified.

Taken together, our data show for the first time that Complex V inhibition in breast cancer cells induces a cell cycle arrest in G0/G1 involving the cell cycle-related kinase AURKA. AURKA was previously found to act on mitochondrial functions (Bertolin et al., 2018; Grant et al., 2018; Bertolin et al., 2021), and we here provide evidence that the kinase can directly interact with the Complex V core subunits ATP5F1A and ATP5F1B. Altering AURKA, ATP5F1A or ATP5F1B levels induces dramatic metabolic changes which affect both the mitochondrial respiratory chain and glycolysis rates. Noteworthy, lowering the abundance of ATP5F1A or ATP5F1B is an efficient way to reproduce G0/G1 arrest in cells with an oxidative metabolism, but it turns on cell death programs in cells with a glycolytic metabolism. Overall, we propose that targeting the AURKA/ATP5F1A/ATP5F1B nexus is a promising avenue to lower breast cancer cell metabolism and reduce cell proliferation.

## Discussion

Our data deepen our understanding of the link between cell cycle progression and mitochondrial functions, with a particular relevance for mitochondrial metabolism. We provide evidence that the G0/G1-phase arrest observable with oligomycin is a direct consequence of Complex V inhibition, as it can be reproduced with the downregulation of the core Complex V subunits *ATP5F1A* and *ATP5F1B* in the absence of mitochondrial insults. Moreover, we show the involvement of the cell cycle machinery in regulating such arrest of the cell cycle. In conditions where the ATP synthase is pharmacologically inhibited, the levels of AURKA dramatically decrease and cells are arrested at G0/G1. Not only AURKA is a cell cycle protein with a clear mitochondrial localization and acting on mitochondrial functions at multiple levels (Bertolin et al., 2021, 2018; Grant et al., 2018), but it also mechanistically links cell cycle progression, Complex V functionality and the cellular metabolic capacity by interacting with Complex V subunits.

In addition to that of AURKA, the abundance of the cell cycle-related protein CCND1 was also lowered upon Complex V inhibition. The lowered expression of CCND1 had previously been linked with metabolic rewiring and enhanced oxidative phosphorylation (Sakamaki et al., 2006), and reductions in CCND1 levels accompanied with G1-phase arrest were observed in HeLa cells after oligomycin treatment (Gemin et al., 2005). However, our data showed that CCND1 is lowered not only upon Complex V inhibition, but also upon Complex III inhibition. This supports the notion that among the cell cycle-related proteins that we screened, AURKA was the only one specifically involved in maintaining Complex V functionality. This role is most likely executed through its physical proximity with ATP5F1A and ATP5F1B. Although CCND1 has never been located at mitochondria so far, this does not exclude its involvement in organelle functions. CCND1 is a protein that orchestrates progression into the G1 phase and G1/S transition in association with CDK4/6 (Tchakarska and Sola, 2020). Prior reports also showed that CCND1 has a functional interaction with AURKA. CCND1 levels lower in conditions where AURKA is overexpressed, and this leads to a failure in undergoing a functional G1/S transition (Jantscher et al., 2011). Therefore, the AURKA/ATP5F1A/ATP5F1B nexus may also include a role of CCND1 in conditions where Complex V is inhibited, although its exact contribution and to what degree its role differs upon Complex III inhibition remains to be defined.

Using FRET/FLIM microscopy, we showed that AURKA is in physical proximity with the ATP5F1A and ATP5F1B core subunits of the mitochondrial ATP synthase. Such proximity is compatible with protein-protein interactions, and it contributes to the maintenance of both mitochondrial and glycolytic ATP production rates. Compromising the integrity of the AURKA/ATP5F1A/ATP5F1B nexus by downregulating any of its components significantly decreases cellular metabolism. Not only the alteration of this nexus is relevant to decrease the rate of ATP produced through the respiratory chain, but it also suggests that glycolysis is unable to act as a compensatory mechanism upon *AURKA, ATP5F1A* or *ATP5F1B* downregulation. These findings raise the exciting hypothesis that a structural alteration of Complex V – by impacting the abundance of any component of the AURKA/ATP5F1A/ATP5F1B nexus – may prevent the ATP synthase to function in reverse mode and maintain ΔΨ (Chevrollier et al., 2011). Although future studies will elucidate the exact functioning of the ATP synthase upon the alteration of the AURKA/ATP5F1A/ATP5F1B nexus, we observed that the consequences of lowered OXPHOS and glycolysis rates are different according to the cell line used and appear to be intrinsic to the cellular metabolic preference. The G1- phase arrest observed upon *AURKA*, *ATP5F1A* or *ATP5F1B* downregulation is specific to oxidative cell lines that express high levels of AURKA, such as MCF7 and T47D cells. Conversely, no impact on the cell cycle was seen in Hs 578T cells expressing low levels of AURKA and relying more on glycolysis. In this model, *AURKA, ATP5F1A* or *ATP5F1B* downregulation rather triggered cell death, possibly implying that the cellular metabolic propensity is a key determinant for cell fate. The metabolic preference of Hs 578T cells towards glucose has previously been reported, since the use of the glucose analog 2-Deoxy-D-Glucose (2-DG) in this cell model lowered cell migration and invasiveness properties (O’Neill et al., 2019). This is further supported by our observation that the abundance of the glycolytic marker protein LDH is higher in Hs 578T cells than in MCF7 and T47D cells. High LDH levels in Hs 578T cells could favor the conversion of pyruvate into lactate, with less pyruvate available to enter the TCA cycle. Last, the preference of Hs 578T cells towards glycolysis was further confirmed with their failure to grow in galactose-containing media.

An exciting, open question that remains to be answered is at what point of the metabolic cascade AURKA should be placed. While it has been shown that nuclear AURKA promotes glycolytic metabolism by upregulating c-Myc (Nguyen et al., 2021), our previous report show an upregulation of mitochondrial Complex IV upon AURKA overexpression (Bertolin et al., 2018). In this light, future studies will elucidate to what extent the organelle-specific isoforms of the kinase can shape the overall metabolic propensity of cells. Last, we here illustrate that the kinase physically interacts with mitochondrial Complex V, indicating that the complexes of the OXPHOS chain are not only functional, but also direct physical interactors of the kinase. Thanks to these proximities within Complex V, AURKA might act as a potential gatekeeper of mitochondrial respiration. Our data also raise the interesting possibility that the absence of AURKA might induce changes in the three-dimensional structure of Complex V, which could be similar to those triggered by *bona fide* pharmacological inhibitors of this mitochondrial complex.

Last, our data suggest that ATP5F1A and ATP5F1B could be promising targets in AURKA-overexpressing cancers. AURKA is a key factor in the pathogenesis of glioblastoma (Nguyen et al., 2021), and increased levels of ATP5F1A and ATP5F1B were observed in glioblastoma models and are accompanied by an upregulation of oxidative metabolism (Xu and Li, 2016). Our results add a critical piece to the puzzle of AURKA-related cancer pathogenesis by providing novel insights in breast cancer cell models of AURKA overexpression. Since AURKA-specific inhibitors show poor efficacy and a high degree of toxicity in clinical trials (Damodaran et al., 2017), the available inhibitors against Complex V functions (Hong and Pedersen, 2008) could be a valuable resource for combinatorial therapies against AURKA and ATP5F1A or ATP5F1B in AURKA-related cancers in the upcoming future.

## Acknowledgments

We thank S. Dutertre and X. Pinson at the Microscopy Rennes Imaging Center (MRic, *Biologie, Santé, Innovation Technologique* - BIOSIT, Rennes, France) for help and assistance with FRET/FLIM experiments. MRic is member of the national infrastructure France-BioImaging, supported by the French National Research Agency (ANR-10-INBS-04). We also thank L. Deleurme and A. Aimé at the Cytometri cell sorting platform (*Biologie, Santé, Innovation Technologique* - BIOSIT, Rennes, France) for extensive help with FACS experiments. We are grateful to B. Fromenty (NuMeCan, Univ. Rennes, France) for help with Seahorse experiments together with S. Rochdi (NuMeCan, Univ. Rennes, France) and Cyrille Berra (IGDR, Univ Rennes, France), and for critical reading of the manuscript together with M. Tramier (IGDR, Univ. Rennes, France). This work was supported by the *Centre National de la Recherche Scientifique* (CNRS), the University of Rennes, the *Ligue Contre le Cancer Comité d’Ille et Vilaine et du Finistère* and the *Association pour la Recherche sur le Cancer* (ARC) to G.B. R.K.S. was supported by a fellowship from *Région Bretagne* (Brittany region, *Stratégie d’Activité Durable – SAD* call, France).

## Author contributions

R.K.S. designed, performed and analyzed the experiments and drafted the manuscript; A.C. performed and analyzed the experiments; G.B. supervised the work, designed the experiments, wrote the manuscript and provided funding.

