## Supplementary Figures for "Aurora kinase A/AURKA interacts with the mitochondrial ATP synthase to regulate energy metabolism and cell death"

### Supplementary Figures and Figure legends

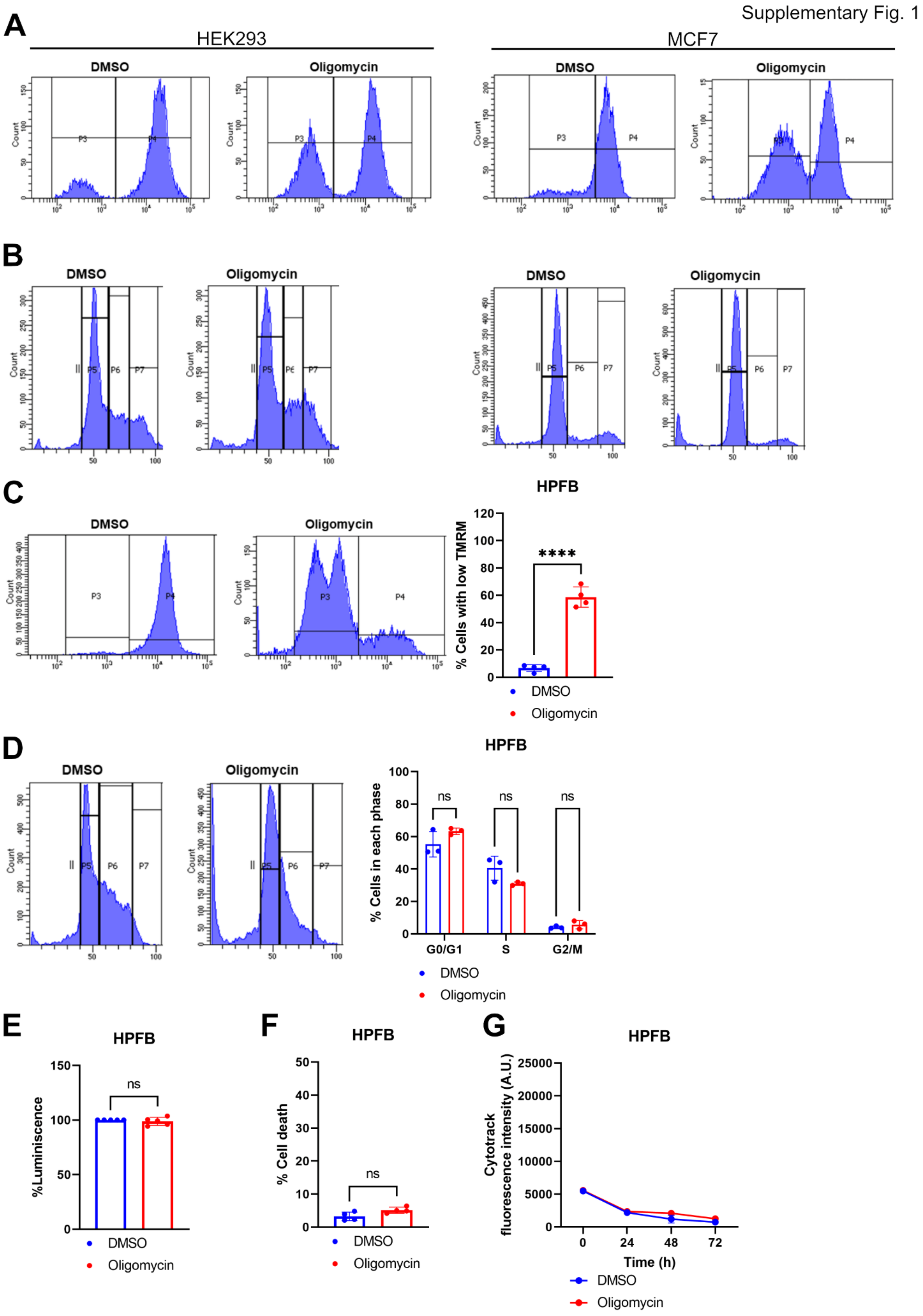

**Supplementary Figure 1. Oligomycin increases the quantity of cells with low TMRM intensity, but it does not trigger cell cycle alterations, ATP loss or cell death in HPFB.** (**A**) FACS plots illustrating TMRM fluorescence intensity in HEK-293 (left) and MCF7 (right) cells treated with DMSO or oligomycin for 48 h. (**B**) FACS plots of HEK-293 (left) or MCF7 (right) cells illustrating Hoechst 33324 intensity to identify the stages of cell cycle, and following an incubation with DMSO or oligomycin for 48 h. (**C, D**) FACS plots and representative quantifications of TMRM intensity (**C**) or Hoechst 33324 incorporation to characterize cell cycle progression (**D**) in HPFB cells treated with DMSO or oligomycin for 48 h. (**E**) Percentage of total ATP levels in HPFB cells treated with DMSO or oligomycin for 48 h. ATP levels were measured using a luminescence-based assay, and were relative to the DMSO condition. (**F**) Percentage of cells showing cell death events and identified with PI/annexin stainings upon treatment with DMSO or oligomycin for 72 h. (**G**) Cytotrack mean fluorescence intensity in HPFB cells treated with DMSO or oligomycin for the indicated time points. Data are means ± SD. ****$P<0.0001$ compared to each corresponding DMSO condition (C-F), or to each corresponding time point in the DMSO condition (G). ns: not significant. $n \geq 3$ independent experiments with at least 10,000 cells per condition quantified.

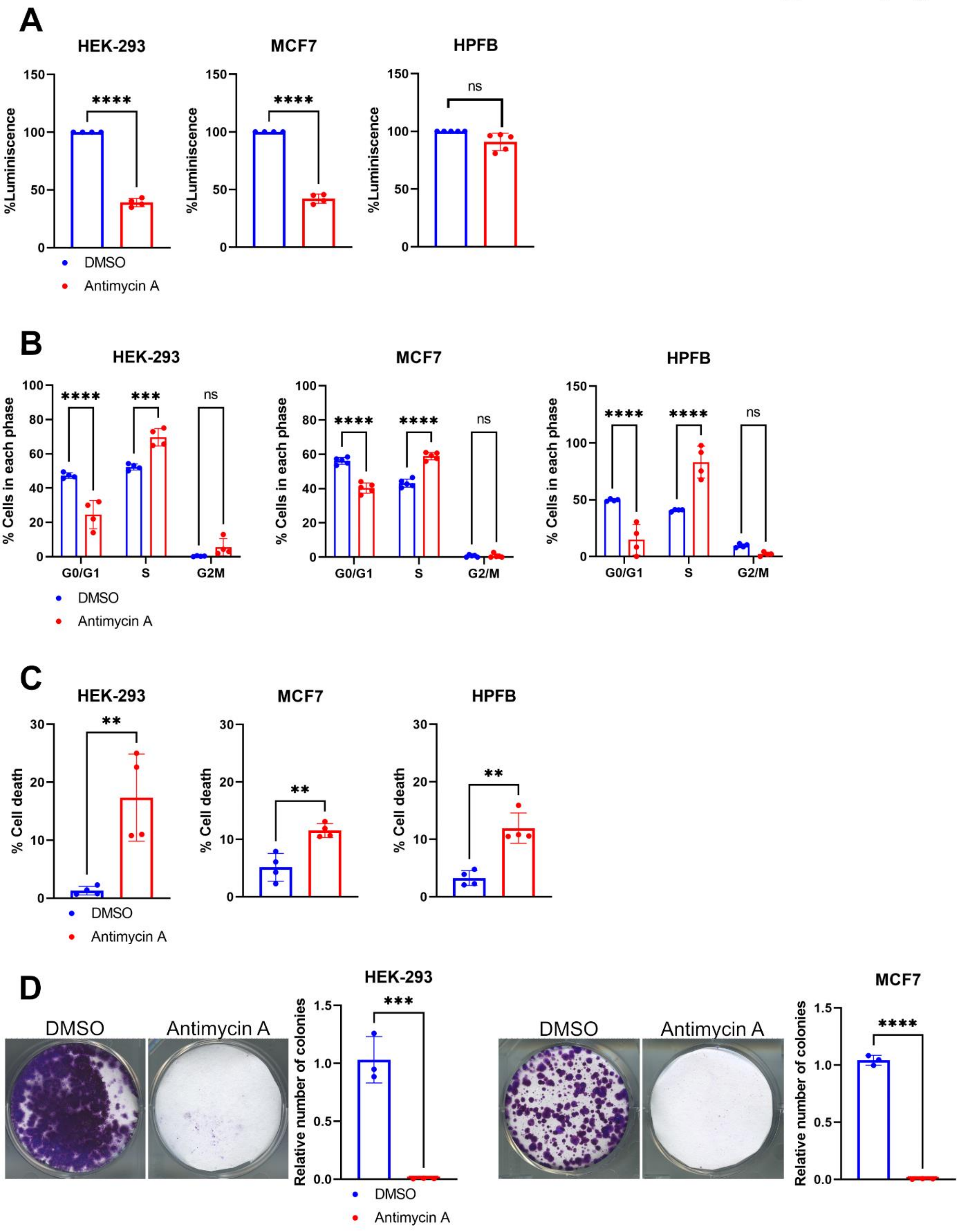


**Supplementary Figure 2. Antimycin A induces ATP loss, S-phase arrest, cell death events and defects in long-term proliferation regardless of the cell model.** (**A-C**) Percentage of total ATP levels (**A**) measured using a luminescence-based assay, and relative to the DMSO condition; quantification of Hoechst 33324 incorporation to characterize the percentage of cells in each cell cycle phase (**B**); percentage of cells showing cell death events and identified with PI/annexin stainings (**C**) in HEK-293 (left), MCF7 (middle), and HPFB (right) cells treated with DMSO or antimycin A for 48 h. (**D**) Representative images and corresponding quantifications of colony-forming assays in HEK-293 (left) or MCF7 (right) cells, treated with DMSO or antimycin A for at least 2 weeks. The number of colonies in the antimycin A -treated condition is relative to that of each corresponding control. Data are means ± SD. $^{**}P<0.01$, $^{***}P<0.001$, and $^{****}P<0.0001$ compared to each corresponding DMSO condition. ns: not significant. $n \geq 3$ independent experiments. At least 10,000 cells per condition were quantified in A-C.

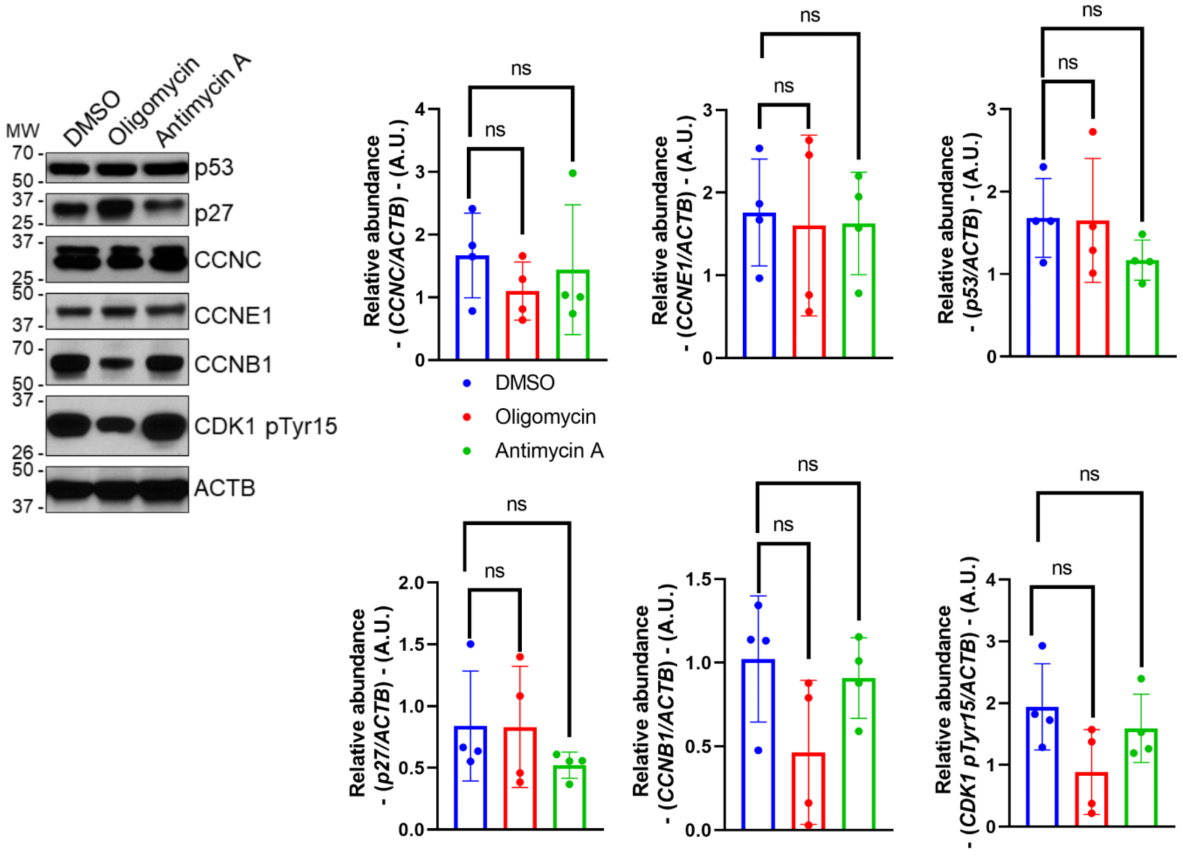


**Supplementary Fig. 3. The abundance of selected cell cycle-related proteins is not lowered upon oligomycin or Antimycin A treatment.** Representative western blot and corresponding quantifications of the abundance of the cell cycle-related proteins p53, p27, CCNC, CCNE1, CCNB1 and CDK1 pTyr15 in total lysates of MCF7 cells treated with oligomycin or Antimycin A for 48 h. Loading control: ACTB. Data are from $n$ = 4 independent experiments, and presented as means ± SD. A. U.: arbitrary units. ns: not significant.

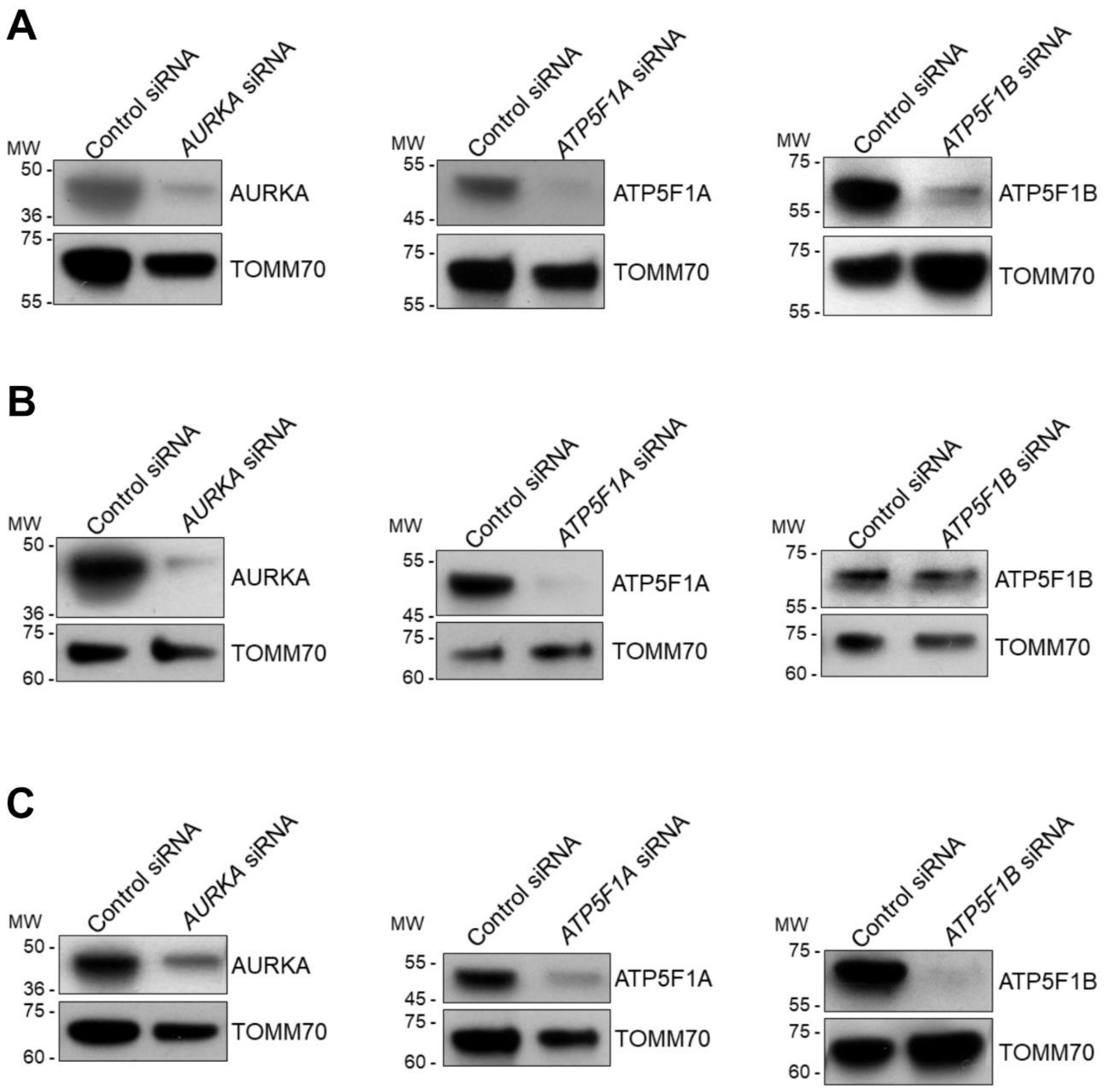


**Supplementary Fig. 4. Knockdown validation strategy of *AURKA*, *ATPF1A* or *ATPF1B*.** (**A-C**) Representative western blots of the abundance of AURKA (left panels), ATP5F1A (middle panels), or ATP5F1B (right panels) in total lysates of MCF7 (**A**), T47D (**B**), or Hs 578T (**C**) cells transfected with a control siRNA or with *AURKA*-, *ATP5F1A*-, or *ATP5F1B*-specific siRNAs as indicated. Loading control: TOMM70.

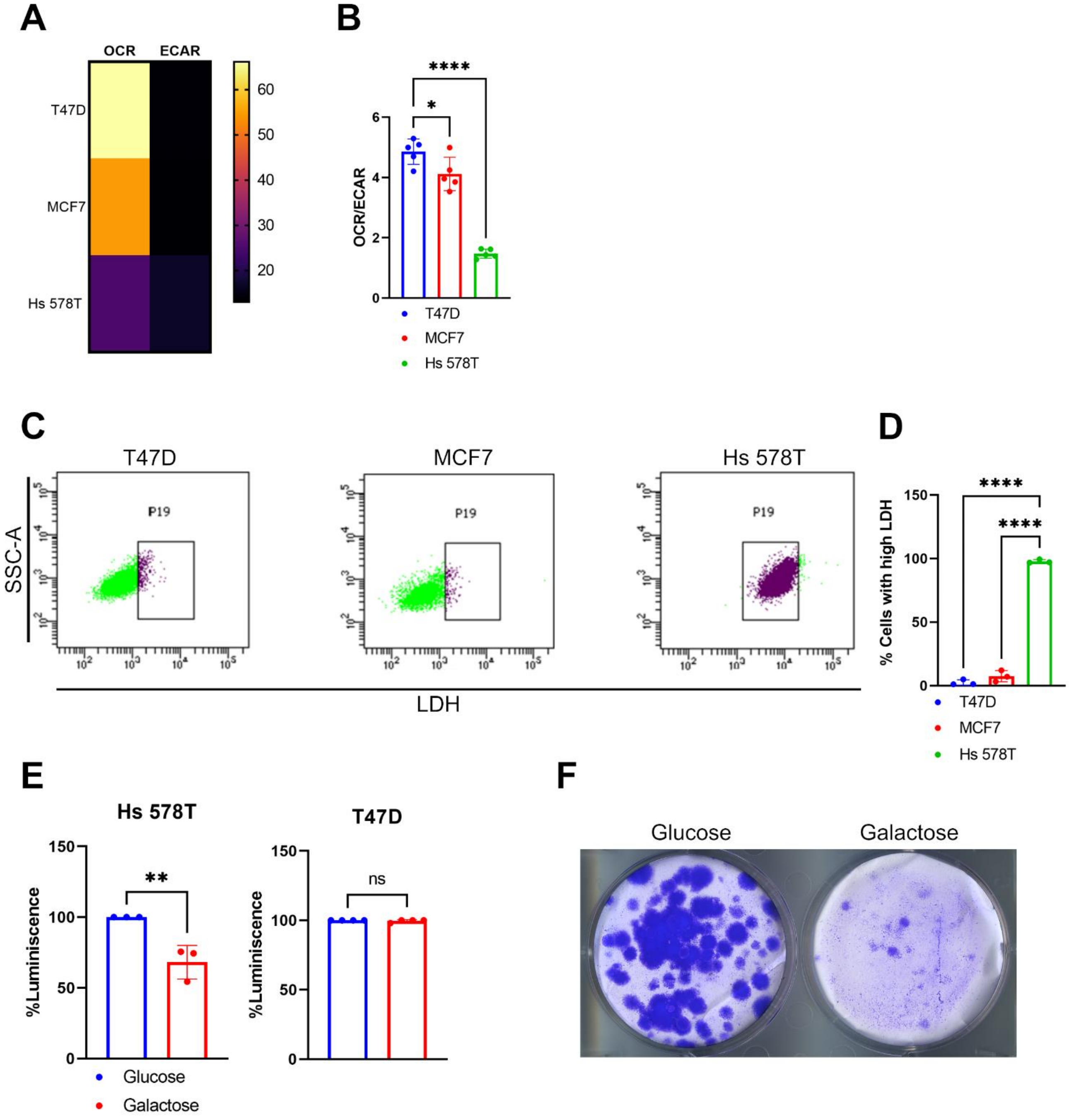


**Supplementary Fig. 5. Hs 578T cells have a glycolytic propensity and fail to proliferate in galactose-containing media.** (**A**) Energy heat map of maximal OCR and ECAR rates from T47D, MCF7 and Hs 578T cells, and derived from Seahorse experiments. (**B**) Maximal OCR/ECAR ratios calculated from basal OCR and ECAR values as in (**A**). (**C**) Representative FACS plots illustrating the LDH-Alexa Fluor 488 staining obtained from T47D (right), MCF7 (middle) and Hs 578T (left) cells. The population of cells showing LDH-specific fluorescence significantly different from background were pseudocolored purple (P19 section). (**D**) Quantification of the percentage of cells in the P19 section in T47D, MCF7 and Hs 578T cells analyzed as in (**C**). (**E**) Percentage of total ATP levels measured using a luminescence-based assay in Hs 578T (left) or T47D (right) cells, grown in media containing glucose or galactose as carbon source. Differences are relative to the glucose condition. (**F**) Representative images of colony-forming assays from Hs 578T cells, grown in media containing glucose or galactose as carbon source for at least 2 weeks. Data are means ± SD. *$P$<0.05,**$P$<0.01 and ****$P$<0.0001 compared to T47D cells (**B, D**) or the glucose condition (**E**). ns: not significant. $n \geq 3$ independent experiments. At least 10,000 cells per condition were quantified in B, D.

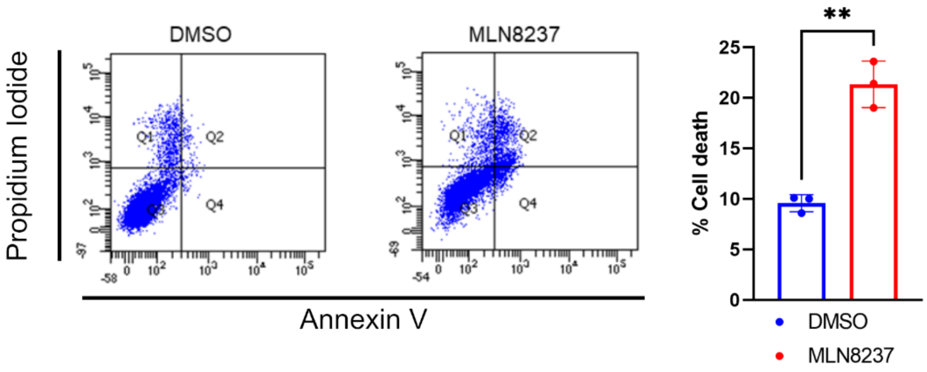


**Supplementary Fig. 6. Hs 278T cells undergo cell death upon AURKA inhibition with MLN8237.** Representative FACS plots and corresponding quantifications (Q2 section) of PI/annexin stainings in Hs 578T cells treated with DMSO or with MLN8237 for 48 h. Data are means ± SD. ***P*<0.01 compared to the DMSO condition. $n \geq 3$ independent experiments. At least 10,000 cells per condition were quantified.
